# Ancestral genomes and population sequencing data reveal strand- and context-specificity of polymorphic G4 sites

**DOI:** 10.1101/2023.09.08.556792

**Authors:** Thamar Jessurun Lobo, Victor Guryev

**Affiliations:** European Research Institute for the Biology of Ageing, University of Groningen, University Medical Centre Groningen, Groningen, Antonius Deusinglaan 1, 9713AV, Groningen, The Netherlands

## Abstract

Recent studies highlight the important functional roles of non-canonical DNA conformations. One of such structures, G-quadruplex (G4), was shown to be involved in multiple processes within a cell such as telomere maintenance, gene regulation, protein translation and alternative splicing. On the other hand, DNA in non-double stranded context can hinder replication and repair processes and, indeed, a higher rate of polymorphisms was reported at G4 sites. However, strand-specificity, positional and nucleotide contexts of nucleotide substitutions at G4 sites are under-investigated.

Here, we combine ancestral genome data and DNA variants from large populational sequencing project to investigate substitution patterns within putative DNA quadruplexes. We confirm the overall elevated rate of base replacement except transitions at CpG sites, which are less likely than in the rest of the genome potentially due to hypomethylated status of G4s. Within G4 loops, there is a trend of replacing weak bases (A, T) with strong bases (G, C) that could promote DNA duplex stabilization. However, this trend is less pronounced when derived allele is rare in the human population. The most over-represented thymine to guanine replacement is about order of magnitude more frequent within G4 than in non-G4 regions. The analysis of nucleotide context of the substitutions shows clear difference between G-rich and C-rich of DNA quadruplexes implying that the strands of the quadruplex might have different mutation dynamics. Finally, we show that the observed deviations from random mutation accumulation result in a biased nucleotide composition of G4 loops rich in adenines. Future studies should reveal specific mutation and selection processes that shape the content of G4-associated DNA polymorphisms.

## Introduction

G4s are secondary structures that can form in single-stranded DNA (or RNA) when at least four stretches of guanines interact through Hoogsteen hydrogen bonds (1). Many years of research have yielded increasing evidence that G4s form in vivo (2). However, consensus on the exact consequences of G4 formation within our cells is still to be reached. On one hand, G4s are suggested to play functional roles in core processes such as telomere maintenance, transcription, RNA splicing and translation. On the other hand, due to non-canonical base pairing, they interfere with DNA replication and thus are associated with genome instability, implying that G4s might not only benefit but also obstruct essential cellular mechanisms such as DNA synthesis (2, 3). Previous studies have investigated germline mutability and selection at G4 loci to help elucidate these apparent contradictory consequences of quadruplex formation. While an early study reported that the single nucleotide polymorphism (SNP) density in G4 sequences was consistently lower (4), this is opposed by more recent papers detailing elevated nucleotide substitution frequencies at G4 loci (5, 6).

Substitution frequencies were especially high in quadruplex sequences that, according to predictions of Quadron software (7), may form particularly stable structures (5). Within G4 sequences, the highest substitution rates were found in the loops, while the guanine stretches contained a significantly higher proportion of SNPs with a lower minor allele frequency (MAF) (5). A follow-up study reports that natural selection maintains a high density and stability of G4 loci in some functional regions of the genome, e.g. 5’ and 3’-UTRs, as well as a low density and stability in others (8). Nonetheless, G4 loci in functional regions including the 5’- and 3’-UTRs were also associated with higher numbers of nucleotide variants (6). Also, splicing-QTLs were found to more frequently overlap G4s that are in close proximity of splice junctions (9).

These observations indicate that G4 structures may indeed perform a critical function in some genomic regions, driving the local preservation of G4-stretches in particular, since they are essential for G4 formation. In addition, the findings suggest that mutation-causing or selection factors affect G4 sequences more strongly, even in genomic locations where they potentially play important roles. Thus, to improve our understanding of how G4 structures may (continue to) influence cellular biology, it seems relevant to know how G4 sequences are changing over time.

In this study we aimed to better characterise DNA base changes that happen in G4 regions. We considered several aspects of mutations at G4 sites that should help us to understand the factors that drive mutations at these regions. First of all, we used the populational frequency as a rough estimate of mutation time of origin. Next to that, we discriminate between ancestral and derived alleles of each mutation by the analysis of a whole-genome alignment of primate species (10). We also take the nucleotide context of the mutation into consideration in order to reveal potential mutation signatures over-represented in G4s. Finally, we considered various types of substitutions in different regions of the DNA quadruplex (G-stretch, middle of the loop or side of the loop) separately. The resulting G4 mutation profiles, observed through these multiple angles, provide insight into the alteration of G4 sequences over time and point to some mechanisms that may be involved in their modification.

## Results

### Most substitution types are significantly increased in G4s

To investigate how single base mutations have affected G4 sequences over time in comparison to non-G4 regions, we determined the associations between common (derived allele frequency, AF >5%), semi-common (between 1% and 5%) and rare SNPs (below 1%) of different nucleotide substitution types and the local presence of a G4 sequence (Figure 1). We inferred the evolutionary direction of mutation at SNPs by employing the human ancestral genome inferred from whole-genome alignments with primate species (10). Because we expected that G4-stretches, the bases of G4-loops adjacent to the G-stretch and the bases in the middle of G4-loops might be affected differently due to their distinct composition and location within the G4 structure, we analysed them separately. Thus, we selected G4 sequences using a G4 motif. G4 sequences were identified by searching the ancestral genome for sequences with four G-stretches consisting of three guanines, each separated by loops of between one and three bases. We only included G4 sequences that were located within experimentally confirmed G4-forming regions (11) of the human genome. We also evaluated CpG and non-CpG dinucleotide contexts separately as the former are known to have a much higher mutation rate.

**Figure 1.**
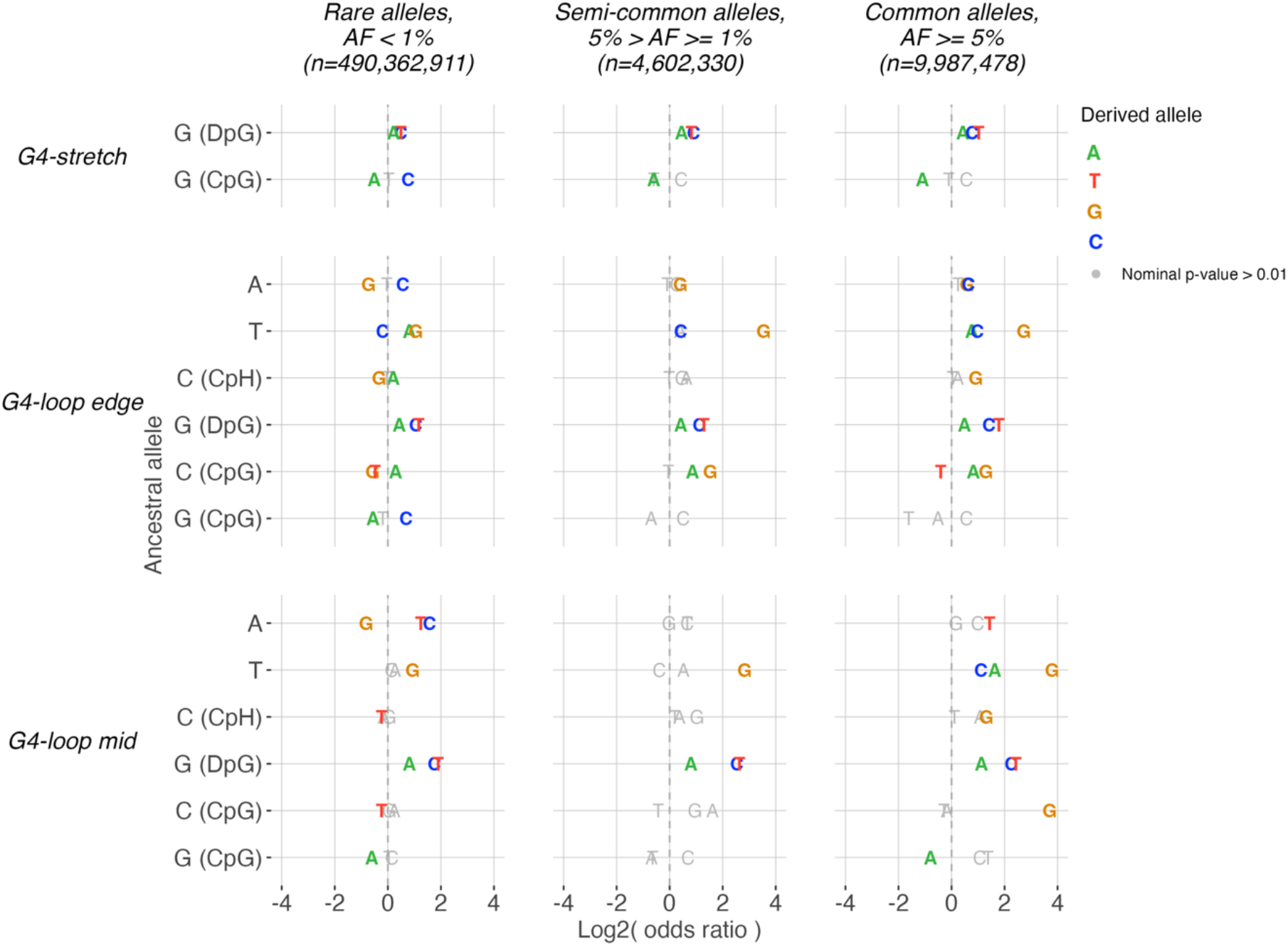
Relative rate of different substitution types in different parts of G4 sequences stratified by derived allele frequencies. A positive log2(odds ratio) indicates that a substitution is more likely to occur within a G4 motif, while a negative log2(odds ratio) indicates that a substitution is less likely to occur within a G4 motif. We determined odds ratios for different sections of the G4 motif independently: the G-stretch, the edge of the loop (single bases located next to a G-stretch) and the middle of the loop. Only associations with a nominal p-value ≤ 0.01 (chi-square test) are displayed in colour, remaining associations are displayed in grey. Associations for substitutions located at CpG and non-CpG sites were plotted separately. Note that the sum of mutations stratified across allele frequencies is not equal to the number of positions per G4 section.

In agreement with the expected high mutability of G4 sequences, we find that most substitution types are more frequently present within G4 motifs than in non-G4 regions (test: Chi-square, nominal p-value < 0.01). This is true throughout different derived allele frequencies and is observed for 19 substitution types seen in rare, 15 in intermediate frequency and 23 in common variants classes. At the same time much fewer substitution types are depleted in G4 regions: 11, 1 and 3 for the aforementioned classes of derived allele frequencies (Figure 1). Particularly guanines that are not in CpG context mutate consistently more often when located within a putative G4 region. As a result, all substitution types that replace ancestral G bases in a non-CpG context are more frequent across different sections of the G4 motif as well as across derived allele frequencies (all nominal p-values ≤ 0.01). We also note that, in general, guanines that are part of quadruplexes mutate more often to cytosines and thymines than to adenines, a trend that is observed across different population frequencies and locations within a G4. In addition, we identify a strong overrepresentation of T-to-G substitutions inside G4-loops. For relatively old polymorphisms from semi-common and common variant groups, the odds ratios vary between 6.6 and 13.9 times.

### Nucleotide transitions are less frequent in CpGs within G4s than in the rest of the genome

Of the substitution types that are rarer in G4 motifs compared to non-G4 regions, most happen at ancestral CpG sites (10 out of 15 with nominal p-values ≤ 0.01). Nearly all of those, nine out of ten, are C-to-T and its reverse complement counterpart G-to-A mutations. In fact, C→T and G→A substitutions occur less frequently at CpG sites in all G4 motif sections for each substitution type (e.g. for rare variants, nominal p-values for all nucleotide transitions in a CpG context are below 0.01). It is well known that spontaneous deamination of methylated cytosines is a common source of transitions at this dinucleotide. Thus, the exclusive depletion of these substitution classes at G4 CpG sites might be connected to local hypomethylation.

### G4 loop bases less frequently change to guanines in rare alleles

Interestingly, at G4-loop positions flanking a G-stretch (G4-loop edge), the relative frequency for several substitution types markedly differs between groups representing common and rare derived alleles. For example, common C→G changes occur more often, whilst there is a relative depletion in rare C→G changes (nominal p-values ≤ 0.01 at both CpG and non-CpG sites). Similarly, while loop edges show an increase of A→G and T→C substitutions in common and intermediate frequency alleles, the same types show the opposite trend when derived alleles are rare (all nominal p-values ≤ 0.01). Hence, an apparent trend that distinguishes groups of common and rare derived alleles is a lower rate of changes leading to guanine bases inside G4 loops. As the appearance of loop guanines might increase the G4 stability by increasing the number of potential G4 folding conformations, this might indicate that DNA changes that lead to more stable G4s have better chances to reach higher frequencies in the human population. Alternatively, these inconsistencies might reflect differences in mutation spectra specific to variants that arose recently (rare) and those that happened in the distant past (common). Yet another explanation of the observed differences might involve the DNA duplex-stabilising effect of the change from a ‘weak’ base (A or T paired by two hydrogen bonds) to a strong base (G or C, three bonds) which might reduce the G4-forming capacity, while increasing processivity of DNA synthesis.

### The relative frequencies at which nucleotide substitutions occur in G4s are context- and strand dependent

The observed differences between CpG and non-CpG contexts imply the importance of the nucleotide context for the rate of G4-specific substitutions. For somatic mutations, it is common to explore base substitution types taking their flanking 5’- and 3’-bases into consideration. This way, one can distinguish among mutational processes that lead to different patterns of nucleotide substitutions in a patient or a genomic region. Embracing a similar approach, we investigated the trinucleotide context dependency of the relative variant frequencies in G4 regions. The complete display of all mutation contexts and derived allele frequencies can be found in the Supplementary materials (Figures S1-S3) and the main results of this analysis are shown in Figure 2. To facilitate the easy interpretation of strand-specific (G4 vs C4, reverse-complement strand of quadruplex) effects, we arranged Figures S1-S3 such that associations for mirroring substitution classes in reverse-complement trinucleotide contexts align horizontally (e.g. TCG→TAG is placed next to CGA→CTA). Thus, we aimed to gain more insight into the mechanisms that could be involved in the mutability/retention of G4 sequences and that differ between G4 and C4 strands of quadruplex sequences.

**Figure 2.**
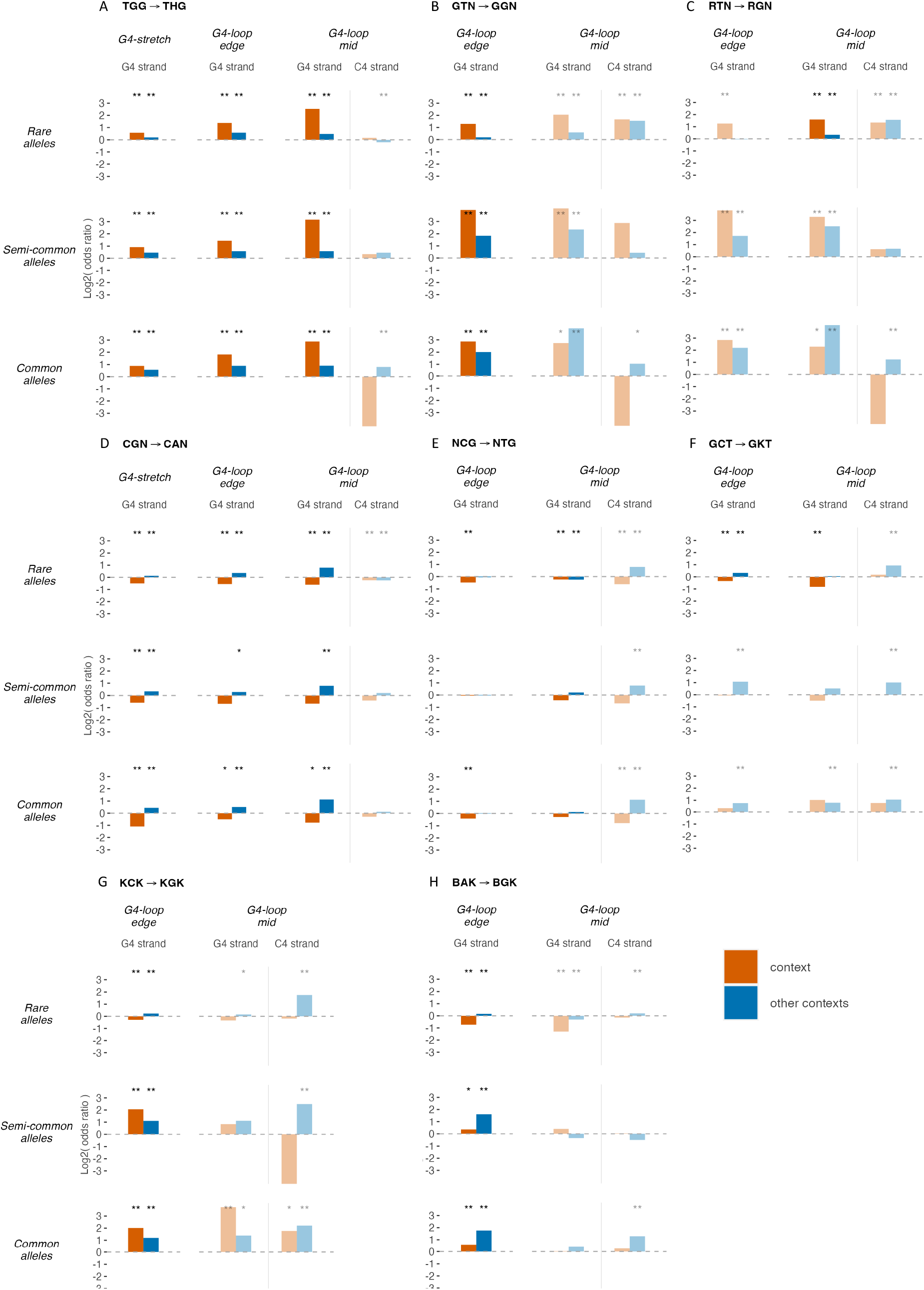
Trinucleotide context-dependent substitution patterns in G4 sequences. We investigated the trinucleotide context dependency of the relative variant frequencies in G4 regions. Panels A-H display eight main patterns, which are indicated at the top of each panel according to the IUPAC codes. The patterns’ odds ratios are indicated by the (highlighted) orange bars, while the (highlighted) blue bars indicate the odds ratios for the same substitution in all other possible/alternative contexts. For example, the highlighted orange bars in panel D indicate that the substitution pattern CGN→CAN is depleted (all log2(odds ratio) < 0) in G4 sequences, while the highlighted blue bars in panel D indicate that DGN→DAN is enriched (all log2(odds ratio) > 0) in G4 sequences. The non-highlighted orange and blue bars indicate the odds ratios for G4 regions or derived allele frequencies that do not display the pattern, or show it to a lesser degree. We also included the odds ratios for the C4 strand at the middle of the loop (CGN→ CAN on the C4 strand at the middle of the loop in case of panel D). Due to the restricted composition of the G4-stretch and the loop edge, odds ratios for the C4 strand of these G4 regions could not be determined (e.g., the GGG trinucleotide context can, per definition, not exist on the C4-strand of the G4-stretch). Nominal significance is indicated by asterisks: * = p-value < 0.05, **= p-value < 0.01. The complete display of variant frequencies in all mutation contexts can be found in the Supplementary materials (Figures S1-S3).

As we have already observed in our previous analysis (Figure 1), non-CpG guanines in G4 sequences mutate more often than guanines in non-G4 sequences. When interpreting these data in the view of the nucleotide context of different substitution types (Figure 2A, Supplementary Figures S1, S2 and S3) we can conclude that this is particularly true for the guanine in a TGG context (all nominal p-values ≤ 0.01, except for G→A in the G4-loop edge and G4-stretch in semi-common and common SNPs). In contrast, cytosine mutations within the reverse complement (CCA) context do not show an overrepresentation within G4 sequences (all nominal p-values > 0.01). This and other examples of discordance between the substitution signatures of the two DNA strands indicate that G4 formation has a profound effect on mutation processes that influence the G4 and C4 strands of quadruplex regions. Further, our analysis of 5’- and 3’-bases highlights guanines in TGG, GGG and AGG contexts within G4 sequences as the ones that are most prone to change to cytosine or thymine, but not to adenine.

### Thymine to guanine substitutions at G4-loops depend on both context and derived allele frequency

At G4-loop edges, T→G replacements with a low derived allele frequency are prominently represented by thymines that have a guanine upstream (Figure 2B, Supplementary Figure S1). Semi-common and common T→G variants are also all more than twice as frequent at thymines with an upstream or downstream guanine compared to non-G4 sites with the same nucleotide build-ups (Figure 2B, Supplementary Figures S2 and S3).

Interestingly, the same substitution is also more prevalent at the C4-strand of quadruplexes. Thus, the complementary substitution type A→C is more frequently seen at the edge of G4-loops for rare and common variants (Figure 1) although it is less pronounced than at the G strand. For rare alleles, A→C substitutions are less context-dependent as, for the significant patterns, the guanine can be found on either side of the substituted base (Figure 2B, Supplemental Figure S1). For semi-common and common alleles, the enrichment of A→C substitutions in the G4-loop edge is not nominally significant for most of the nucleotide contexts (Figure 2B).

At the middle of G4-loops, the higher occurrence of T→G substitutions (for rare alleles) primarily originates from cases where thymines are preceded by an adenine or guanine (Figure 2C, Supplementary Figure S1). Likewise, complementary A→C substitutions in the middle of G4-loops also, mostly, show a higher prevalence within a RAN context.

We also observed a strong increase in semi-common and common T→G replacements in the middle of G4-loops (Figure 1). However, these increases are not nominally significant within most of the trinucleotide contexts (Figure 2C, Supplementary Figures S2 and S3). The latter can be explained by the very limited number of occurrences of the semi-common and common T→G SNPs within those nucleotide contexts.

### No clear trinucleotide context dependency for transition underrepresentation at CpG sites

Figure 1 showed that the substitutions associated with DNA methylation, C→T and G→A, occur less frequently at CpG sites in G4 sequences. For rare G→A substitutions (Figure 2D), this is true for any context containing a CpG site (all nominal p-values ≤ 0.01) within any G4-sequence section (Supplementary Figure S1). Rare C→T changes (Figure 2E), too, are relatively depleted within G4-sequences in every possible CpG site-containing context. However, the associations are not nominally significant for substitutions in GCG and TCG contexts in the middle of G4-loops. Semi-common and common C→T and G→A replacements are not significantly depleted with all CpG siteslog containing contexts in G4 sequences, either (Figure 2D, E; Supplementary Figures S2 and S3). Nonetheless, all nominally significant depletions seen for variants of semi-common and common allele frequencies concern C→T or G→A substitutions at CpG sites.

### Relative depletion of low-frequency substitutions in GCT context

The consequent nominally significant depletion of C→G and C→T changes in rare alleles within GCT contexts of G4-loops (both the middle and the edge of the loop) comprise another noticeable pattern (Figure 2F, Supplementary Figure S1). However, the ‘mirroring’ substitutions G→C and G→A at the reverse complemented AGC context (only present in the middle of G4-loops) are neither always depleted nor nominally significant. We do not observe this pattern for semi-common and common C→G and C→T substitutions.

### Inconsistent enrichment pattern between rare and common derived alleles

Within G4-loop edges, while rare C→G derived alleles are depleted within GCG, GCT and TCG contexts, common ones within ACG, GCG, GCT and TCG contexts do show a relative enrichment (nominal p-values ≤ 0.01, Figure 2G, Supplementary Figures S1 and S3). On the other hand, all nominally significant rare and common complementary substitutions indicate enrichment.

For semi-common C→G substitutions, most contexts do not reach nominal significance except for the GCG triplet, which exhibits an enrichment (Supplementary Figure S2).

Similarly, while we see a depletion of rare A→G variants within GAT, TAG, GAG and CAG contexts at the edge of G4-loops, semi-common and common substitutions of this type are enriched in respectively GAG, AAG, GAA and all possible contexts except for GAC (nominal p-values ≤ 0.01, Figure 2H, Supplementary Figures S1, S2 and S3).

For the ‘mirroring’ substitution, T→C, rare derived alleles are depleted from G4-loops within almost all possible contexts (except for GTT and GTC) at the edges of G4-loops as well. The semi-common and common T→C substitutions at the edges of G4-loops do not reach nominal significance for any of the nucleotide contexts, except for GTT, where it shows enrichment.

### Low rate of transitions at G4 CpG nucleotides can be explained by hypomethylation

As mentioned above, we found that G→A and C→T substitutions occur consistently less frequently at CpG sites located within G4 motifs. This dinucleotide is seen as hypermutable from multiple recent sequencing projects and even reaches saturation (judged by the number of reciprocal independent variants observed at a site) when a large cohort is being sequenced (12). In agreement with this observation, our analysis of GnomAD variants showed 74.7% of ancestral CpG sites had a transition in this database, a strong contrast with 11.7% seen for non-CpG sites. As these transitions are thought to primarily originate from deamination of methylated cytosine bases, the depletion of such substitutions within G4 regions suggests hypomethylation of their GpGs. To test this hypothesis, we employed public methylation profiling data obtained by whole-genome bisulfite sequencing in several human tissues.

The results show significantly lower levels of CpG methylation at both strands (G4 and C4) and parts (loop and stretch) of putative DNA quadruplexes. These results are in line with previous observations associating G4s with a lower DNA methylation status (13). Across the tissues, the C-stretch consistently exhibits lower methylation levels than the quadruplex loop at the G4-strand, suggesting that apart from a prerequisite for double-stranded DNA, cytosine methylation can be shaped by additional mechanisms asymmetrically acting on G4 and C4 strands of DNA quadruplexes.

Alternatively, methylation at CpG sites may occur at different rates depending on the broader nucleotide context, e.g CCCpG versus CpGGG.

### Context and dynamics of G4 creation and elimination in the genomes

Our observation of different mutation dynamics between G4 and non-G4 regions in the genomes prompted us to investigate whether such biases affect the rates of G4 motif formation and elimination from genomes. We considered all uninterrupted ancestral segments of 15 bp and longer and ancestral-to-derived allele substitutions to see which of them result in the creation of a G4 motif and which remove a putative G4 present in the ancestral DNA sequence. To estimate a background distribution, we performed the same analysis with permutation of the substitutions retaining the relative abundance of mutations in each trinucleotide context. The comparison of observed and expected (under random distribution of mutations) can provide us with an interesting insight into the evolution of G4 loci.

### Rates of creation and elimination of G4s differ depending on derived allele frequency

Since G4s may be functionally important, mutations at G4 sites, including those creating or disrupting the quadruplex, could be subject to selection processes. We find that common alleles share a similar number of substitutions that invoke a G4 motif as those that potentially destroy it (Table 1). This, however, is dramatically different for alleles that reached intermediate frequency, where more than two thirds of variants changing G4 capacity are spawning a new G4 motif. On the contrary, over three quarters of rare alleles altering local G4 status eliminate them. Throughout all categories we observed an excess of the number of observed mutations over the number of expected ones (upon permutation). The discrepancy between the two rates differs dramatically from 34% increase (rare alleles eliminating a G4 motif) to 13.7 times difference (intermediate frequency alleles creating a G4 motif).

**Table 1.** Number of derived alleles creating and eliminating G4 motifs, stratified by derived allele frequency.

| Derived allele frequency | Alleles creating G4 motif |  | Alleles eliminating G4 motif |  |
| --- | --- | --- | --- | --- |
|  | Observed | Expected | Observed | Expected |
| Rare (<1%) | 83,485 | 37,926 | 226,195 | 168,524 |
| Intermediate (1-5%) | 4,615 | 336 | 2,204 | 1,526 |
| Common (>5%) | 5,373 | 1,021 | 5,850 | 3,831 |

The dynamics of mutations that lead to the gain or loss of G4 motifs suggest that G4 sequences are indeed more mutable. This can be judged from the excess of rare derived alleles that eliminate a G4. A peculiar observation is that there is an even higher excess of rare alleles that lead to a G4 sequence, suggesting that potential G4 regions might have an elevated mutability even before acquiring a recognizable G4 motif. The apparent differences seen among substitutions with different allele frequencies suggest that variants changing the G4 properties of a region might be under selective pressure. Thus, a more moderate increase of rare variants removing a G4 over a much stronger increase in alleles creating a G4 (each related to their expected background levels, Table 1) might partially be explained by a combination of a higher mutation rate and purifying selection sweeping G4 loss mutations. At the same time, a remarkable abundance of derived alleles with intermediate frequency that introduce a G4 motif (4,615 observed vs 336 expected) might indicate their potential selective advantage combined with slow propagation throughout the population (e.g. under balancing selection).

### Difference in nucleotide context of variants creating or eliminating a G4

When considering the nucleotide contexts of mutations that create G4 motifs, several interesting observations can be made (Figure 4A). First of all, every nucleotide context shows an over-representation of variants compared to the genome-wide rate for the same nucleotide contexts. We identified three groups of contexts that have dramatically elevated levels of variants leading to the creation of a G4 motif. The first group constitutes sites flanked by Gs from both sides (depicted in orange), such as the GTG motif (observed: 24,197, expected: 5,613 variants). The other two groups with exceptionally high odds for G4-forming correspond to T-to-G substitutions with either upstream (blue) or downstream (red) guanine. Note that from all motifs where the substituted nucleotide is bracketed by Gs (GAG, GCG, GTG), the one that corresponds to T-to-G replacement shows the highest odds for creating G4 motifs.

**Figure 3.**
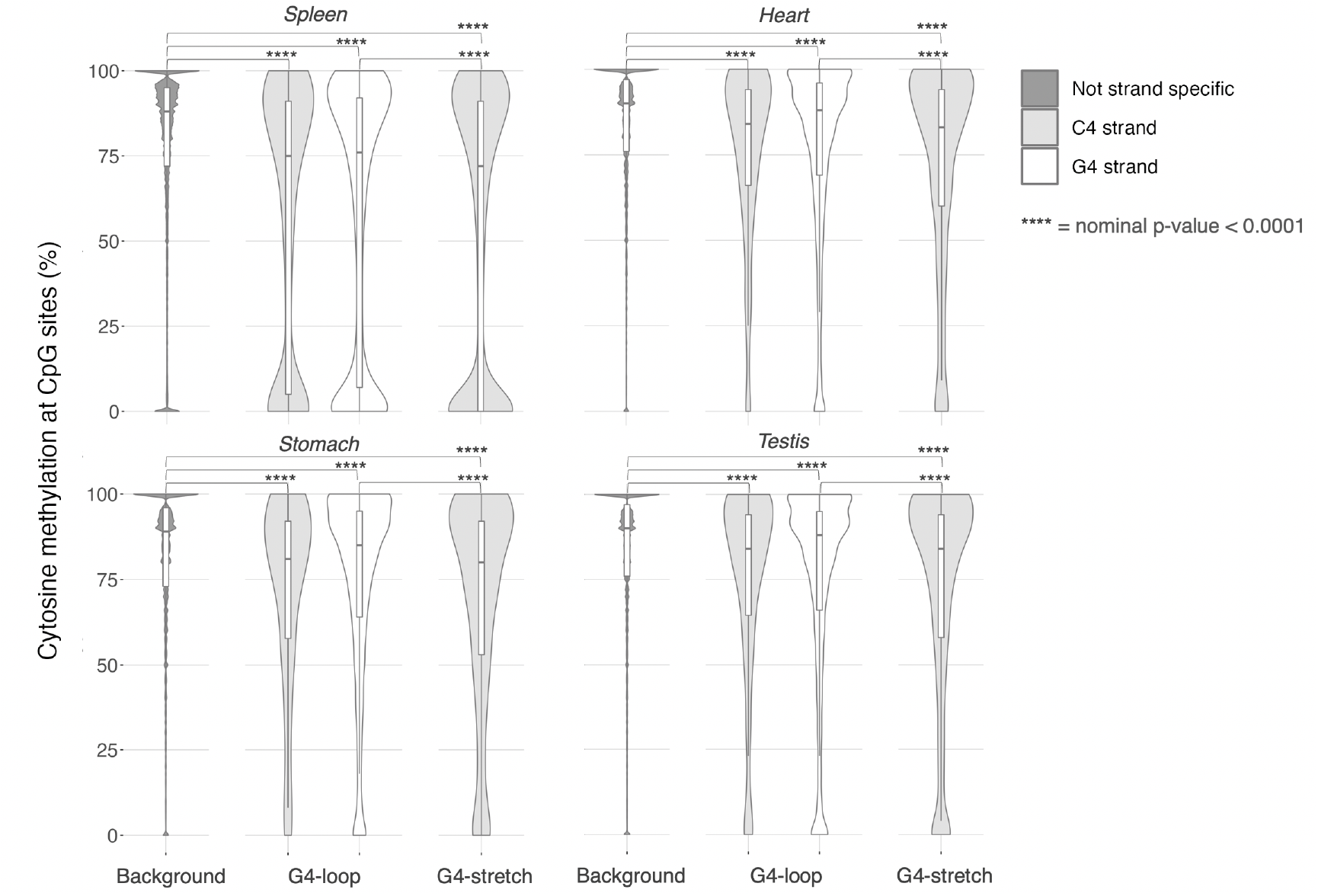
WGBS data show that CpG sites in G4 sequences are hypomethylated. Distributions of cytosine methylation percentages at CpG sites in tissue from human spleen, heart, stomach and testis differ significantly between G4 sequences and background (all nominal p-values < 0.0001). Median methylation percentages are lower in G4-loops and G4-stretches. Exact nominal and adjusted p-values can be found in Supplementary Table S1.

**Figure 4.**
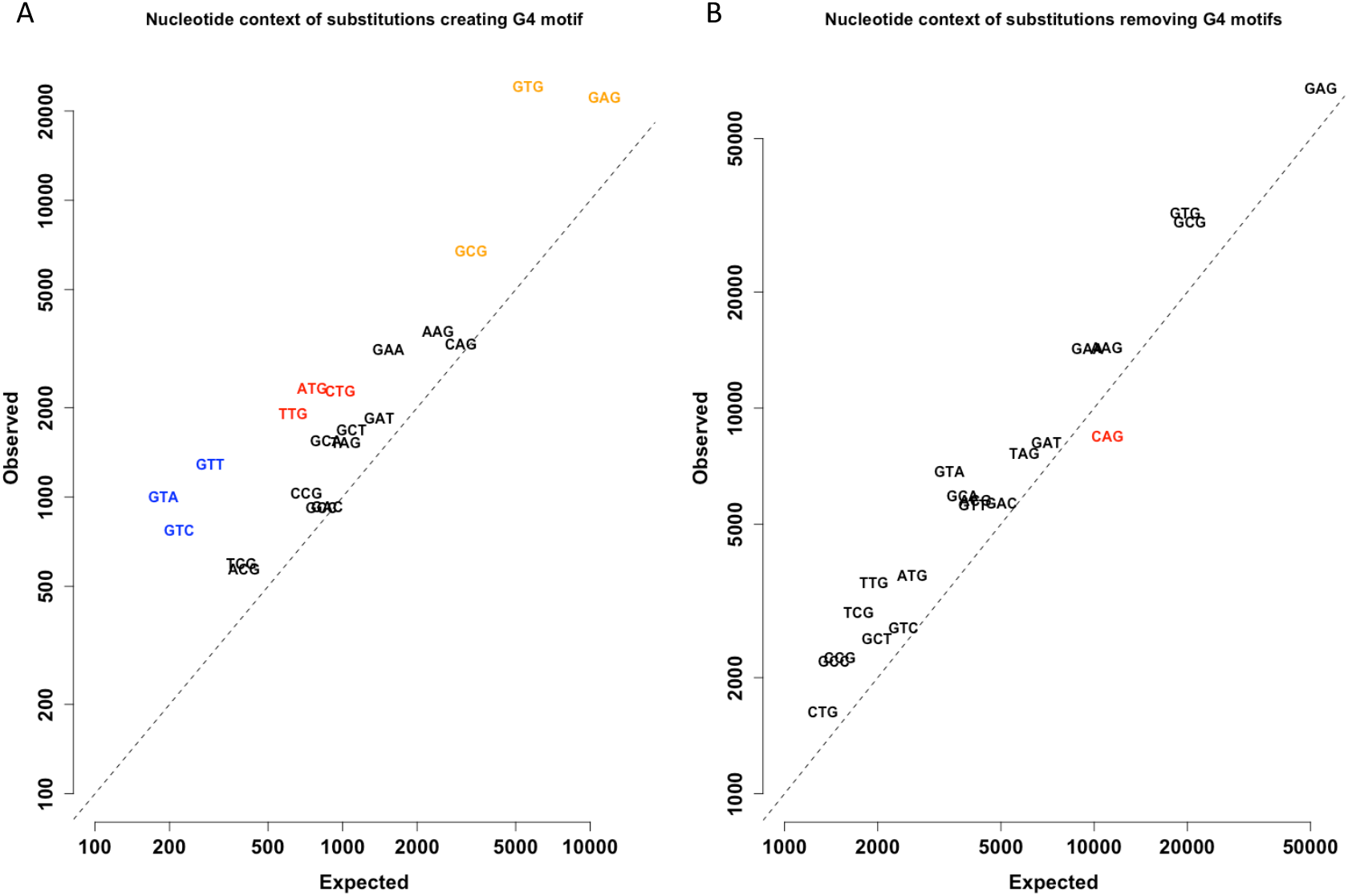
Substitutions creating or eliminating G4 motifs have different occurrences that depend on their nucleotide context. A. Number of observed and expected (permutation analysis) variants creating G4 motifs. Ancestral trinucleotides are displayed for which substitution of the middle base to G leads to creation of a G4 motif. Motifs that have guanines bracketing substitution sites are given in orange, while motifs where T→G substitutions lead to a G4 are highlighted with blue (upstream G) and red (downstream G). The diagonal line represents an equal amount of observed and expected substitutions, both axes are log-scale. B. Number of observed and expected (permutation analysis) variants eliminating G4 motifs. Derived trinucleotides are displayed for which substitution of the middle guanine base leads to loss of a G4 motif. The motif resulting from a transition at CpG sites (CGG→CAG) is highlighted in red. The diagonal line represents an equal amount of observed and expected substitutions, both axes are log-scale.

Similarly, most of the nucleotide contexts where derived alleles disrupt a G4 motif show increased levels of DNA variants compared to an average genomic region (Figure 4B). The notable exception is CGG→CAG change (shown in red), which is the only context that corresponds to the only option of transition at CpG dinucleotide that can result in G4 motif removal. These observations are compatible with a generally higher mutability of G4 sites as well as relative hypomethylation of CpG sites within putative G4 regions.

### Rates of G4-creating and eliminating variants depend on derived allele frequency

When stratifying substitutions by the frequency of derived allele we observed considerable heterogeneity between groups (Figure 5). Overall, biases characteristic to substitutions resulting in the appearance of a new G4 motif are much more pronounced for derived alleles that are common or semi-common in the population. Previously observed trends of increase of T-to-G changes flanked by upstream and/or downstream guanines and a stronger enrichment of variants with an intermediate frequency of the derived allele are clearly observed as well. We do not observe such extreme biases among variants that remove G4 motifs. We note a milder depletion of CpG related G4 disrupting variants in rare compared to common or intermediate groups. On the contrary, several contexts/substitution combinations show a higher prevalence in (semi-)common rather than in rare derived alleles (e.g. TGG→TTG or GGA→GAA).

**Figure 5.**
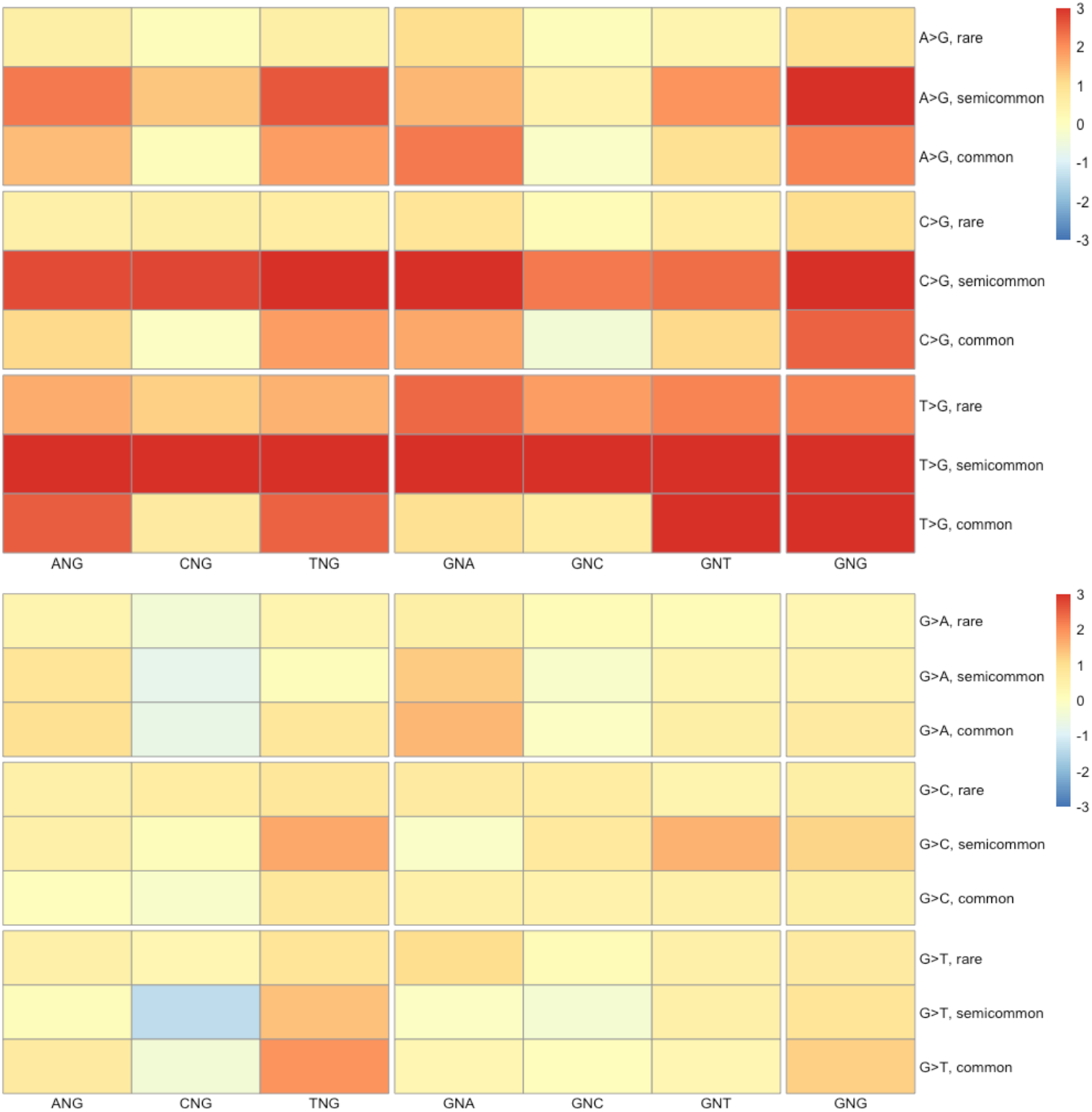
Relative occurrence of variants creating G4 (top panel) and removing G4 (bottom panel) for different nucleotide contexts, substitution types and frequencies. Log2 ratio of observed vs expected number of variants per nucleotide context (columns), substitution type (row, groups) and different derived allele frequencies (rows, within groups).

### Biases in G4-related substitution patterns affect G4 composition in human genome

The observed disproportions in abundance of different substitution types that drive the gain and loss of G4 sequences should have consequences for the genome-wide composition of G4 motifs. We therefore investigated which sequence context of G4-forming regions occurs more frequently in our genome. While such abundance of a particular type of G4 region might result from mutation bias and selection pressures, it can also stem from repeat propagation, e.g. when a G4 sequence is an integral part of a mobile element. The list of G4 sequences that are frequently observed in non-repetitive parts of the human reference shows that they are strongly biased (Table 2). Apparently, commonly found motifs typically have short loops enriched in adenines. While, upon random nucleotide distribution in a genome with a lower GC context (such as human’s), we would expect a similar number of G4s where all loops are single adenosines to those where all loops are single thymidines, the actual numbers are quite different (2913 vs 324 respectively). Apart from the latter, simple motif, the only two motifs that do not have exclusive adenosine-G4-loops are motifs where most of the genomic occurrences are due to being part of the L1PA or L1HS retrotransposons. It is likely that many of their non-repetitive entries descend from more ancient LINE1 integrations and thus are not independent events.

**Table 2.**
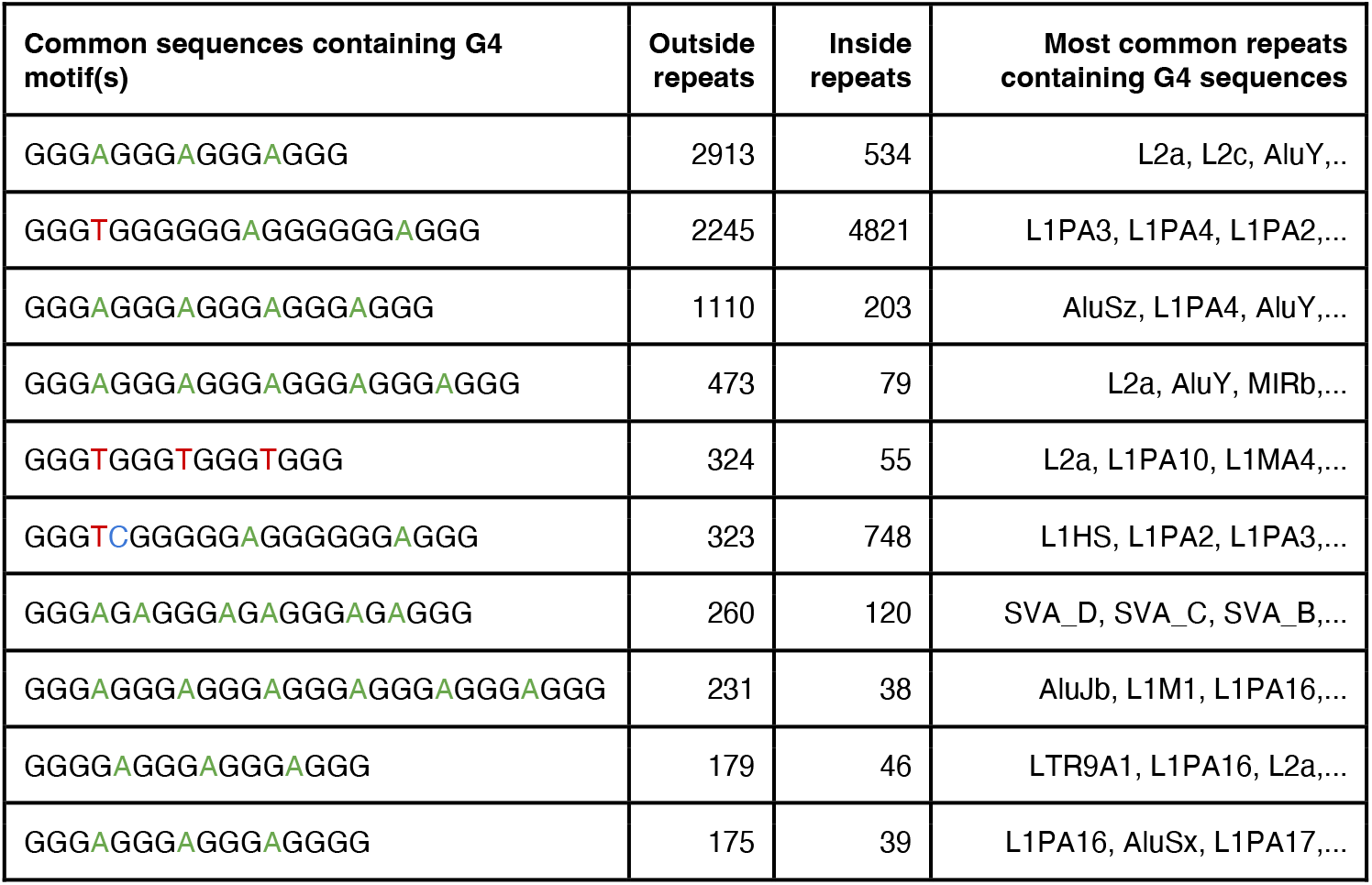
Ten most frequent G4 motif-containing sequences in the non-repetitive part of GRCh38 genome reference. Number of G4 regions (concatenating overlapping G4 motifs with loop sizes 1-3bp). Regions that are located completely inside an annotated repeat (except simple, satellite, unknown repeats or low complexity regions) are counted separately.

The observed trend for T→G changes, especially when flanked by guanine bases upstream and/or downstream can lead to three anticipated consequences: i) removal of thymidines from the G4 loops; ii) shortening of the loops and iii) expansion of the G-stretches. Our analysis of the most common putative G4-forming sequences in the human genome reference indeed reflects the presence of these trends and supports non-random dynamics shaping contemporary genome-wide quadruplex composition.

## Discussion

It has been observed by previous studies that putative G4 quadruplex genomic regions have a higher density of DNA variants, implicating them as fast-evolving parts of our genome (5, 6). However, questions about which substitution types preferentially happen, whether G-stretches and G4-loops, C4 and G4 quadruplex strand show differences in this respect and whether these processes affect rare and common alleles similarly remained largely unanswered.

In order to explore these questions, one needs to know the ancestral and derived state of each polymorphism and use genome sequences in its ancestral state for prediction of mutation directionality. In this study, we employed an ancestral genome reconstructed from primate whole-genome alignment (10) and combined it with experimental data on G4 identification, G4 motif prediction methods and a large catalogue of genomic variants (14).

While our results confirm the higher rate of substitutions at putative quadruplex sites, we also make some interesting observations about the origin and context of these DNA changes. Although the vast majority of substitution types increase their abundance at G4s compared to the non-G4 regions, C-to-T transitions do not seem to follow this rule. This trend is especially strong for cytosines within a CpG context, indicating a potential connection to a specific DNA methylation pattern of G4s themselves.

Indeed, existing whole-genome methylation data across several tissues shows that G4 regions are hypomethylated and thus should be less prone to such transitions. Interestingly, we observed slight, but significant differences between the methylation profiles of quadruplex loops at the G4 strand and G-stretches at the C4 strand (C-stretch), hinting that the low methylation status is not solely a result of a G4’s capacity to avoid methylation by staying outside of the classical double-stranded DNA form. However, even outside of the CpG context we rarely observed a significant increase of C-to-T transitions throughout G4 parts and derived allele frequencies. On the contrary, changes that lead to ‘strong’ bases - Gs and Cs seem to be more abundant within putative G4s. This might be a strategy to ‘stabilize’ the double-stranded DNA conformation at G4 sites.

It is worth noting that the latter pattern is more clearly seen among derived alleles that are common or have an intermediate frequency in the human population (at least 1% in gnomAD database). Although most of the over-representations of substitution types seem consistent among groups with different derived allele frequencies, the differences indicate that either a preference for mutations was stronger in the past (when many of the common alleles originated) or that this noticeable difference is due to large-scale selective processes that drive allele fixation at G4 sites. Future studies should investigate the population structure and variant dynamics to reveal the relative importance of these two processes on the increased variation within G4 loci.

The difference in biases between the variants located within DNA quadruplexes and those in the rest of the genome might stem from deficient repair processes outside of the double-stranded DNA context. It was therefore interesting to see if the same substitution within the same nucleotide context will have a different probability depending on its localization on the G- or C-strand of the G4 quadruplex. As we observed multiple cases where the G4-forming strand has a different bias than its complementary strand, we conclude that vulnerability and/or DNA repair act differently for G- and C-strands of the DNA quadruplex. Our current analysis was unable to implicate either of the strands as being more prone to harbour new mutations. This question should be addressed in future studies using techniques that map sites of DNA damage in a strand-specific manner.

The biases observed in this study prompted us to investigate the process of gain and loss of G4 motifs by the introduction of new alleles. It is peculiar that non-G4 regions that acquire a G4 motif through the introduction of a new derived allele, show an enrichment in mutation rates for all nucleotide contexts compared to the rest of the genome (Figure 4A). That means that either pre-G4 regions might already possess an elevated mutation rate or that mutations that lead to G4s are more frequently retained in the population. Further, the gain and loss of G4s only shows an equilibrium for common alleles, while intermediate frequency (1-5%) alleles are twice as common to give rise to a G4 than to remove it. On the contrary, rare, likely recently acquired alleles are 2.7 times more likely to remove a G4 motif rather than create it.

Finally, the preferential patterns of DNA changes, such as T-to-G mutations for thymidines flanked by guanines, impose that G4 loop size and length themselves might be biased through this directional accumulation of derived alleles. Our survey for G4 motifs within the non-repetitive part of human genome reference indeed shows that most frequently observed loops are short and predominantly contain adenines rather than thymines. Also here, we need to determine whether this is a result of the G4-specific mutation spectrum or can be an adaptive feature that drives differential retention of alleles corresponding to A-rich short-loop G4s in the population.

## Materials and Methods

### G-quadruplex selection

Locations of experimentally defined G4 forming DNA sequences in the human genome in K^+^ conditions were downloaded from GEO accession GSE110582 in BED format. Coordinates were lifted from hg19 to hg38 assembly using the UCSC lift-over Genome Annotations tool with default settings. We used a custom Perl script to select sequences with the G4 motif G[3]N[1-3]G[3]N[1-3]G[3]N[1-3]G[3] (n= 389,698; 87,417 regions when overlapping sequences are merged) within these known G4-forming genomic regions in the autosomal human GRCh38 ancestor genome (http://ftp.ensembl.org/pub/release-106/fasta/ancestral_alleles/homo_sapiens_ancestor_GRCh38.tar.gz). Within the resulting set of sequences, we defined G4-stretches as G[3] (n bases = 829,104) and G4-loops as N[1-3] (n bases = 406,331), where we discriminate between edge-loop positions (located right next to a G4-stretch) and mid-loop positions. We excluded base positions that, regarding the used G4 motif, could be marked as both part of a G4-stretch and a G4-loop, and base positions in non-uniquely mappable genomic regions (selected based on hg38/k100.umap.bed.gz).

As background, we used the human autosomal GRCh38 ancestor genome minus non-uniquely mappable regions, all G4-forming regions found with G4-seq and any occurrences of our G4 motif from the ancestral genome (n bases = 2,502,697,538).

### Variant selection

We downloaded single base variants (gnomad.genomes.v3.1.2.sites.vcf) from gnomAD v3.1.2 (https://gnomad.broadinstitute.org/) (12). We selected autosomal single base variants with a pass filter. To ensure that the reference allele reflected the ancestral state in the human GRCh38 ancestor genome, we excluded variants where neither reference nor alternative allele was equal to the ancestral base and exchanged reference and alternative alleles for variants where the alternative allele was equal to the ancestral base. In the latter case, we adjusted the allele frequency (AF) accordingly. Variants in non-uniquely mappable genomic regions were excluded. Finally, we categorized each variant as common (AF ≥ 5%, n=490,362,911), semi-common (5% > AF ≥ 1%, n=4,602,330) or rare (1% > AF, n=360,012).

### Mutation rate quantification

For each substitution class, we quantified its number of occurrences in G4 loops, G4 stretches and background in a context aware manner (meaning including the bases immediately 5’ and 3’ of the ancestral genome). When a variant was located inside a G4 on the reverse strand, we reverse complemented it. We also quantified the total number of occurrences of each 3-base combination, again, in a strand aware manner in case of a G4. Log2(odds ratios) for each substitution class were calculated both in a tri-nucleotide context-aware and -ignorant manner per G4 section. P-values were calculated using a Chi Square test with Monte Carlo simulation through the chisq.test function in R version 4.2.1. Plots were created with R package ggplot2 (version 3.3.6).

### Analysis of WGBS datasets to compare CpG C-methylation in G4s to background

We selected four human Whole-Genome Bisulfite Sequencing (WGBS) datasets from the Encyclopedia of DNA Elements (ENCODE) consortium (https://www.encodeproject.org/) and compared CpG cytosine methylation percentages between G4 stretches, G4 loops and the background. Samples for the selected datasets were taken from spleen (https://www.encodeproject.org/files/ENCFF336UWV/), heart (https://www.encodeproject.org/files/ENCFF318IKY/), testis (https://www.encodeproject.org/files/ENCFF926RAG/), and stomach (https://www.encodeproject.org/files/ENCFF586VVJ/). We downloaded processed bed bed9+ files with the methylation states at CpG sites from ENCODE4 v1.1.5. We categorized autosomal methylation states based on location in a G4-stretch, G4-loop or background (see *G-quadruplex selection* section) and strand (G4-strand, C4-strand or both strands in case of the background). We excluded Cs with coverage below 10 reads. Violin plots of the distributions were created using the R package ggplot2 (version 3.3.6). We did not plot distributions for C-methylation states at G4-stretches on the G4-strand, as G4-stretches are supposed to only contain Gs on the G4-strand. Cs found in those locations are a consequence of mismatches between the ancestral human genome we use for G4 selection and the human reference genome GRCh38 used for WGBS mapping. The equality of the median CpG methylation percentages between the categories was tested with a Kruskal-Wallis rank sum test followed by a Dunn’s test of multiple comparisons with the Benjamini Hochberg method in R version 4.2.1.

### Nucleotide context analysis of derived alleles in G4

We parsed the human ancestral genome sequence for blocks that are equal or longer than 15bp (and thus can harbour a G4 motif). We only analysed bases that had an upstream and downstream base for which ancestral context was known. We required that all three bases should be mappable by NGS sequencing (that is overlap with regions from k100 mapping track from UCSC genome browser). Next, we overlapped these positions with coordinates of biallelic base substitutions from GnomAD database (version 3.1.2) taking into account allele frequency in population (AF flag in VCF file). Derived allele frequency was calculated as allele frequency of alternative allele if reference base was matching ancestral base or as allele frequency of reference allele if ancestral base was matching the alternative allele. The number of each substitution type and each ancestral triplet (upstream, replaced and downstream bases) were recorded. G4 motifs were predicted using aforementioned patterns with allowed loop sizes between 1 and 3 base pairs. Overlapping G4 motifs were clustered in G4 regions. Within G4 region bases were assigned to the G4 loop if they none of the G4 motifs they were part of G4-stretch, otherwise they were treated as part of G4 stretches. Triplets and nucleotide context for G4s on negative strands were reverse-complimented in order to discriminate between biases that are different between G-rich and C-rich strands of DNA quadruplexes. Expected number of mutations was estimated using simulation of random mutations across the same ancestral blocks while keeping the chance and allele frequency of mutations specific to each ancestral triplet.

### Analysis of derived alleles that result in gain or loss of G4 motif

For each ancestral block as defined above (including size and NGS mappability requirements) we fetched biallelic SNV variants from GnomAD. For each variant we modified the ancestral base to derived allele and run G4 motif predictions for original and modified versions. The cases where G4 motifs were overlapping the substituted base only in derived sequence were classified as G4 gain alleles, while cases where G4 motifs were seen at that position only in original/ancestral block were called as G4 loss alleles. Location within G4-loop/stretch, G4 strandedness, derived allele frequencies and nucleotide contexts (triplets) were taken into account, as described above.

### Analysis of common G4 motifs in GRCh38 genome reference

The prediction of G4 motifs was done as described above, including merging overlapping G4 motifs, but excluding procedure for retaining NGS-mappable regions. We clustered G4 regions based on their primary nucleotide sequence and counted occurrence of each in the human genome reference. We also noted the number of the sequences that are located inside regions annotated as repeats (e.g. retrotransposons, or DNA transposons) that can arise in our genomes through the genomes by ‘copy-pasting’ or ‘cut-and-pasting’ mechanisms and not through base mutation processes. For this reason, we did not consider simple repeats, low complexity regions or unknown repeats as they are likely acquiring a G4 motif via base substitution process.

## Data availability

The code generated during this study is available at GitHub; https://github.com/thamarlobo/G4_mutation_analysis.

## Supporting information

**Figure S1.**
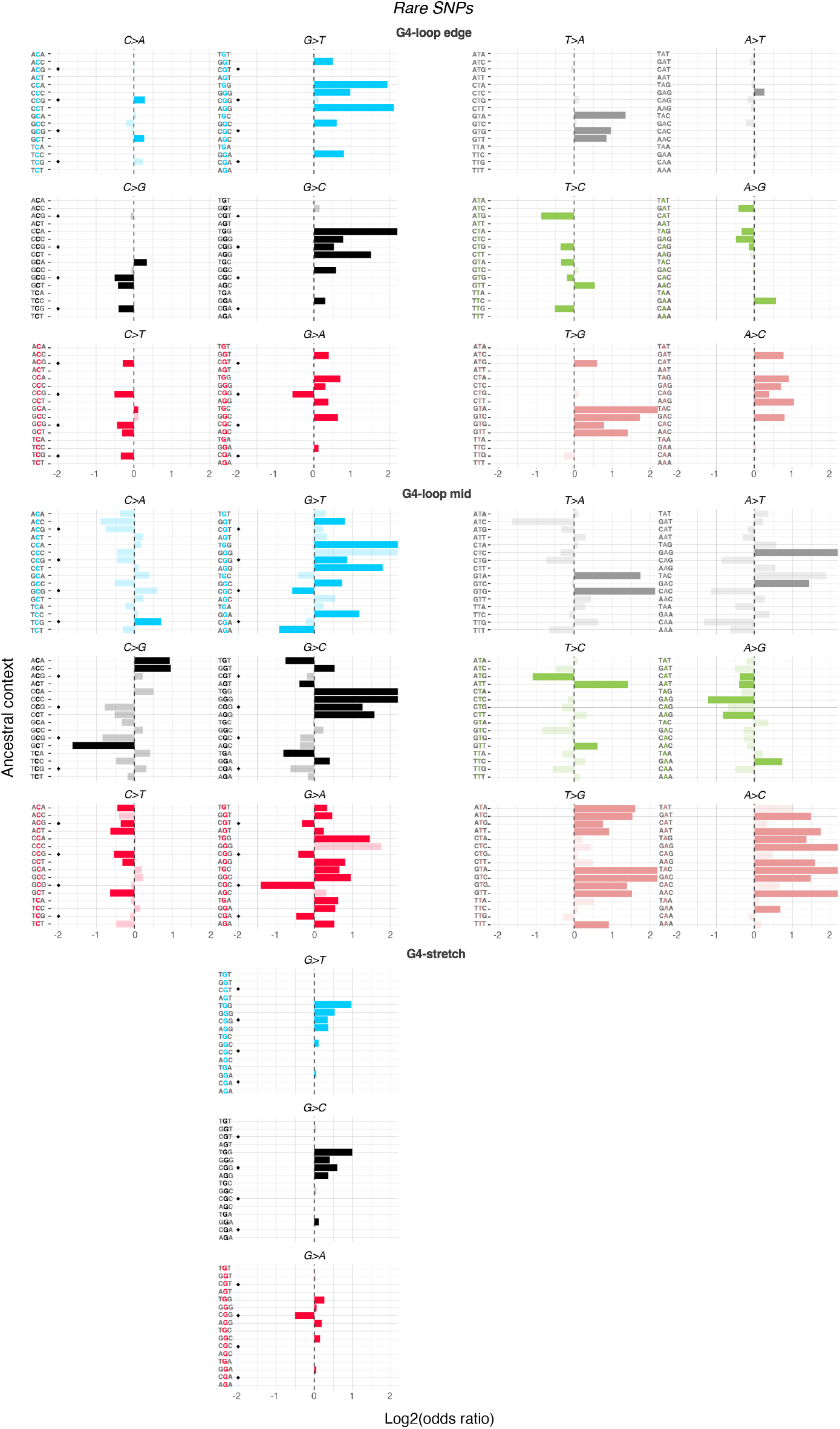
Associations between rare SNPs of different nucleotide substitution classes and the presence of a local G4 motif per ancestral context. A positive log2(odds ratio) indicates that the substitution in the specified context is more likely to occur within a G4 motif, while a negative log2(odds ratio) indicates that the substitution in the specified context is less likely to occur within a G4 motif. We determined odds ratios for different sections of the G4 motif independently: the G-stretch, the edge of the loop (single bases located next to a G-stretch) and the middle of the loop. Semi-transparency indicates a nominal p-value > 0.01 (Chi-square test). Contexts that include a CpG site are marked by a black diamond at log2(odd ratio) = −2. Note that, within the G4-loop edge, at least one of the flanking bases must be a G, and within the G4-stretch, the mutated base and at least one of the flanking bases must be a G. To facilitate the easy interpretation of strand-specific (G4 vs C4, reverse-complement strand of quadruplex) effects, we arranged the figure such that associations for mirroring substitution classes in reverse-complement trinucleotide contexts align horizontally.

**Figure S2.**
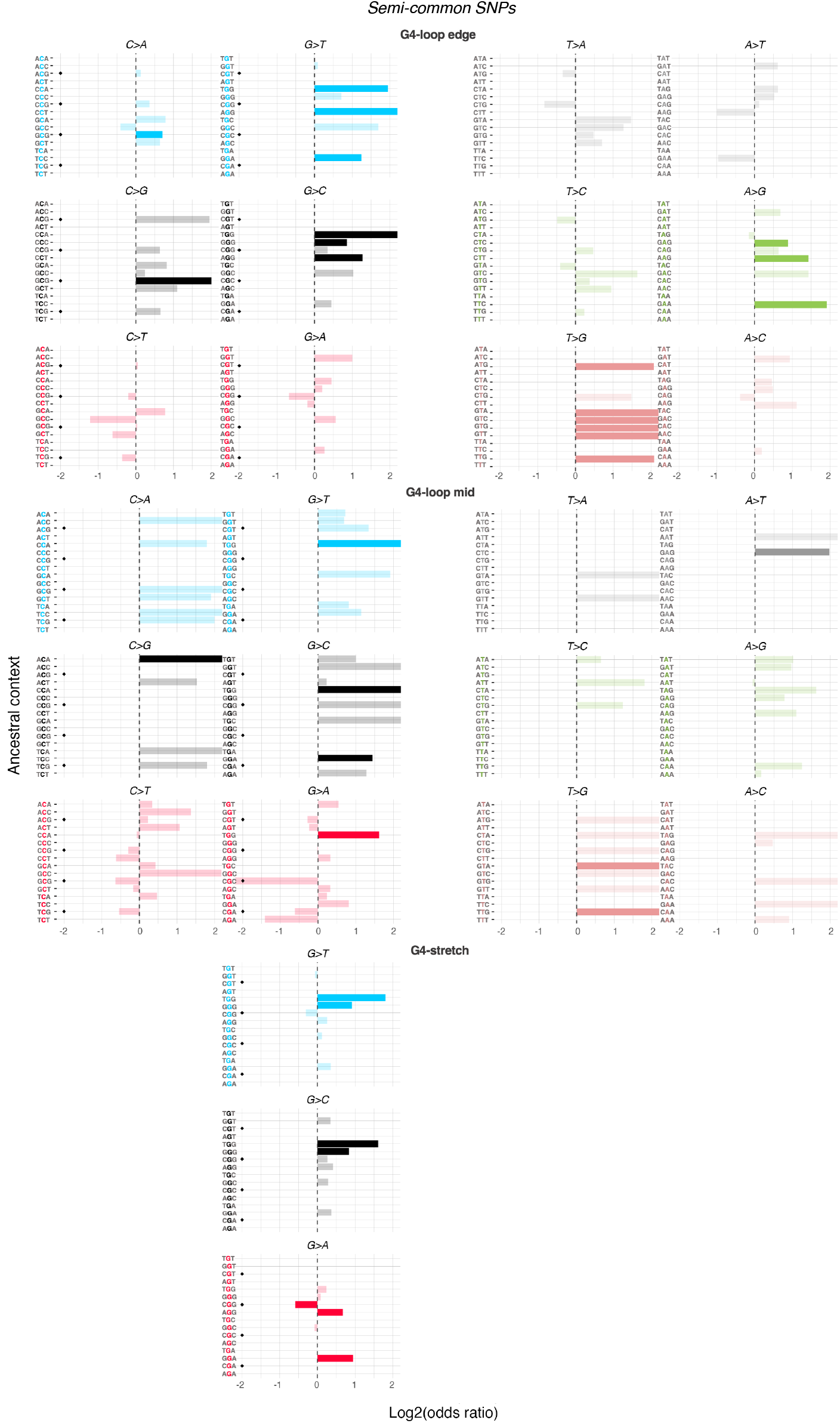
Associations between semi-common SNPs of different nucleotide substitution classes and the presence of a local G4 motif per ancestral context. A positive log2(odds ratio) indicates that the substitution in the specified context is more likely to occur within a G4 motif, while a negative log2(odds ratio) indicates that the substitution in the specified context is less likely to occur within a G4 motif. We determined odds ratios for different sections of the G4 motif independently: the G-stretch, the edge of the loop (single bases located next to a G-stretch) and the middle of the loop. Semi-transparency indicates a nominal p-value > 0.01 (Chi-square test). Contexts that include a CpG site are marked by a black diamond at log2(odd ratio) = −2. Note that, within the G4-loop edge, at least one of the flanking bases must be a G, and within the G4-stretch, the mutated base and at least one of the flanking bases must be a G. To facilitate the easy interpretation of strand-specific (G4 vs C4, reverse-complement strand of quadruplex) effects, we arranged the figure such that associations for mirroring substitution classes in reverse-complement trinucleotide contexts align horizontally.

**Figure S3.**
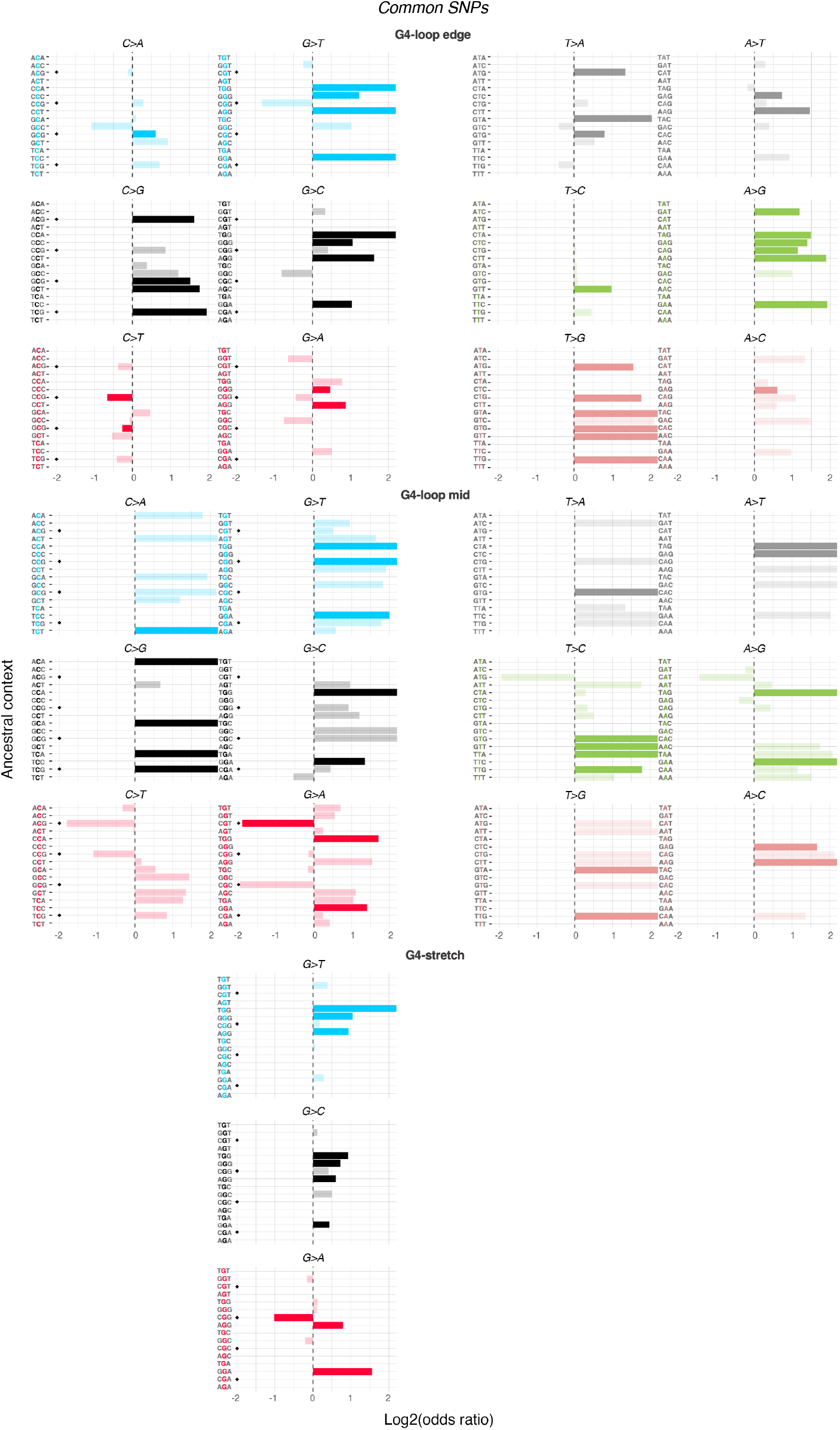
Associations between common SNPs of different nucleotide substitution classes and the presence of a local G4 motif per ancestral context. A positive log2(odds ratio) indicates that the substitution in the specified context is more likely to occur within a G4 motif, while a negative log2(odds ratio) indicates that the substitution in the specified context is less likely to occur within a G4 motif. We determined odds ratios for different sections of the G4 motif independently: the G-stretch, the edge of the loop (single bases located next to a G-stretch) and the middle of the loop. Semi-transparency indicates a nominal p-value > 0.01 (Chi-square test). Contexts that include a CpG site are marked by a black diamond at log2(odd ratio) = −2. Note that, within the G4-loop edge, at least one of the flanking bases must be a G, and within the G4-stretch, the mutated base and at least one of the flanking bases must be a G. To facilitate the easy interpretation of strand-specific (G4 vs C4, reverse-complement strand of quadruplex) effects, we arranged the figure such that associations for mirroring substitution classes in reverse-complement trinucleotide contexts align horizontally.

**Table S1.** Nominal and BH adjusted p-values from Dunn’s Post Hoc test comparing CpG site C-methylation percentage distributions between G4-loops, G4-stretches and background for WGBS datasets from human spleen, heart, stomach and testis tissue.

| Tissue \| Comparison | Z | P unadjusted | P adjusted (Benjamini-Hochberg) |
| --- | --- | --- | --- |
| **Spleen** |  |  |  |
| G4-loop ( C4 strand ) - G4-stretch | 2.208049 | 2.724085e-02 | 3.27E-02 |
| G4-loop ( C4 strand ) - G4-loop ( G4 strand ) | -1.611186 | 1.071393e-01 | 1.07E-01 |
| G4-stretch - G4-loop ( G4 strand ) | -7.566739 | 3.827087e-14 | 5.74E-14 |
| G4-loop ( C4 strand ) - Background | -21.814810 | 1.68E-105 | 3.36E-105 |
| G4-stretch - Background | -55.927983 | p < 2.225074e-308 | p < 2.225074e-308 |
| G4-loop ( G4 strand ) - Background | -61.733687 | p < 2.225074e-308 | p < 2.225074e-308 |
| **Heart** |  |  |  |
| G4-loop ( C4 strand ) - G4-stretch | 0.597561 | 5.50E-01 | 5.50E-01 |
| G4-loop ( C4 strand ) - G4-loop ( G4 strand ) | -1.778 | 7.54E-02 | 9.05E-02 |
| G4-stretch - G4-loop ( G4 strand ) | -5.07726 | 3.83E-07 | 7.66E-07 |
| G4-loop ( C4 strand ) - Background | -4.11951 | 3.80E-05 | 5.70E-05 |
| G4-stretch - Background | -10.8132 | 2.98E-27 | 8.95E-27 |
| G4-loop ( G4 strand ) - Background | -11.327 | 9.64E-30 | 5.79E-29 |
| **Stomach** |  |  |  |
| G4-loop ( C4 strand ) - G4-stretch | 0.371608 | 7.10E-01 | 7.10E-01 |
| G4-loop ( C4 strand ) - G4-loop ( G4 strand ) | -2.42674 | 1.52E-02 | 1.83E-02 |
| G4-stretch - G4-loop ( G4 strand ) | -6.00841 | 1.87E-09 | 3.75E-09 |
| G4-loop ( C4 strand ) - Background | -4.99748 | 5.81E-07 | 8.71E-07 |
| G4-stretch - Background | -12.3574 | 4.44E-35 | 1.33E-34 |
| G4-loop ( G4 strand ) - Background | -12.8926 | 4.95E-38 | 2.97E-37 |
| **Testis** |  |  |  |
| G4-loop ( C4 strand ) - G4-stretch | 1.058815 | 2.90E-01 | 2.90E-01 |
| G4-loop ( C4 strand ) - G4-loop ( G4 strand ) | -1.9487 | 5.13E-02 | 6.16E-02 |
| G4-stretch - G4-loop ( G4 strand ) | -6.04248 | 1.52E-09 | 2.28E-09 |
| G4-loop ( C4 strand ) - Background | -6.170384 | 6.81E-10 | 1.36E-09 |
| G4-stretch - Background | -16.0698 | 4.16E-58 | 2.49E-57 |
| G4-loop ( G4 strand ) - Background | -15.8966 | 6.69E-57 | 2.01E-56 |

